# The *RAD51* paralogue *HvXRCC2* regulates meiosis and recombination in barley

**DOI:** 10.1101/2023.10.05.561044

**Authors:** Isabelle Colas, Malcolm Macaulay, Mikel Arrieta, Miriam Schreiber, Jamie Orr, Amritpal Sandhu, Anu Augustine, Małgorzata Targońska-Karasek, Susan J Armstrong, Robbie Waugh, Luke Ramsay

## Abstract

Using a positional candidate-gene approach we show that semi-sterile *desynaptic8* mutants are associated with deletions in or complete knockout of the barley homolog of *XRCC2* (*X-Ray Repair Cross Complementing 2*). In barley *XRCC2* mutants, the initial meiotic progression is normal, albeit with a small delay in initiation, with completion of synapsis. However, the absence of *HvXRCC2* subsequently leads to a dramatic reduction in the number of crossovers, chromosome mis-segregation, and infertility, suggesting that *HvXRCC2* plays a major role in recombination. This mutant phenotype is congruent with that reported in mammalian studies but contrasts with the *XRCC2* mutant in Arabidopsis which is fertile, exhibits normal chromosome pairing and correct chromosome segregation, and is associated with an increased rate of crossovers. This indicates that the *XRCC2* mutant phenotype in Arabidopsis is not representative of all plants and that *XRCC2* is not a good candidate for the modulation of recombination in barley.

**Highlight:** The mutants of the barley homolog of *XRCC2* exhibit delays in replication leading to defective meiosis, altered RAD51 orthologue behaviour, and significant reduction in the number of crossovers as in canonical mammalian *XRCC2* mutants but unlike those in Arabidopsis.

## Introduction

Meiosis is the specialized process required for sexual reproduction that involves two cell divisions after a single round of DNA replication that results in gametes with half the genetic complement of the parent (Alberts, 2015). During meiosis I, two major interlinked events occur simultaneously: chromosome synapsis and homologous recombination (HR). Synapsis aligns the homologous chromosomes along their entire length, providing the necessary proximity to promote effective recombination (Zickler, 2006). Homologous recombination (HR) is essential for chromosome segregation during the first division of meiosis but also for the exchange of genetic material via crossovers that create unique allelic combinations in each of the four haploid gametes (Wang and Copenhaver, 2018; Zickler, 2006). During meiosis, HR starts with SPO11 catalysed DNA double strand breaks (DSB) which are subsequently resected to generate a 3’ single stranded DNA (ssDNA) end (reviewed in Mercier et al., 2014). The 3’ ends are then captured by the RecA-like proteins RAD51 and DMC1 (meiosis specific) to promote strand invasion and DSB repair by the homologous DNA template (Brown and Bishop, 2014). During meiosis, this strand invasion is essential for the formation of a “D-loop” which leads to a crossover when repaired via the double Holliday junction (dHJ) pathway (Daley et al., 2014; Wang and Copenhaver, 2018).

Five *RAD51* paralogues (*RAD51B*, *RAD51C*, *RAD51D*, *XRCC2,* and *XRCC3*) have been found in both mammals and plants (Da Ines et al., 2012; Da Ines et al., 2013a; Sullivan and Bernstein, 2018). These form two distinct complexes: BCDX2 (RAD51B-RAD51C-RAD51D-XRCC2) and CX3 (RAD51C-XRCC3), which are both involved in meiotic and somatic recombination, DNA repair, and chromosome stability (Sullivan and Bernstein, 2018; Chun et al., 2013; Masson et al., 2001).

Effective null mutation of *XRCC2* leads to a delay in (but not an absence of) RAD51 foci formation in hamster cells (Liu, 2002) and numerous studies have shown that various *XRCC2* mutants are hypersensitive to DNA cross-linking agents such as mitomycin C (MMC), cisplatin, aldehydes, tirapazamine and temozolomide and that DNA replication fork dynamics are disturbed (Liu, 2002; Liu et al., 1998; Liu and Lim, 2005; Saxena et al., 2018; Wang et al., 2014). In Arabidopsis and more recently in Drosophila, *XRCC2* has also been shown to be important for somatic recombination and DNA repair under genotoxic stress while having a minor or no evident role during meiosis (Bayer et al., 2020; Bleuyard et al., 2005; Da Ines et al., 2013a; Wang et al., 2014). This contrasts with mammalian studies where *xrcc2* effective null mutations cause chromosome mis-segregation (Cui et al., 1999; Griffin et al., 2000; Mozdarani et al., 2001), developmental defects (Adam et al., 2007) and meiotic arrest and infertility (Griffin et al., 2000). Infertility also results from a point mutation (41T>C) in human and mouse *XRCC2* (Yang et al., 2018). Female Drosophila *xrcc2* mutants on the other hand are fertile with normal development, which has been attributed to a postulated redundancy of the proteins XRCC2 and XRCC3 (Bayer et al., 2020). In Arabidopsis, *xrcc3* null mutants are defective for meiosis, but *xrcc2* null mutants are fertile and exhibit normal chromosome pairing, synapsis, and correct chromosome segregation (Bleuyard et al., 2005; Bleuyard and White, 2004; Da Ines et al., 2013a). Interestingly, the *xrcc2* null mutant in Arabidopsis was also reported to be associated with an increased rate of crossovers and recombination (Da Ines et al., 2013a).

There is considerable interest in mutants in genes such as *FANCM*, *RecQ4,* and *Hei10* which increase crossover number in Arabidopsis due to their potential application in crop breeding (Mieulet et al., 2018; Arrieta et al., 2021). The potential to modulate recombination is particularly attractive in large genome cereals such as barley, where crossover distribution patterns are strongly skewed toward the ends of the chromosomes, limiting the potential recombination of large sections of the genome (Kunzel and Waugh, 2002; Lambing and Heckmann, 2018; Mascher et al., 2017). The behaviour of *XRCC2* mutants in plant species other than Arabidopsis is thus of interest given potential breeding and genetic applications.

*desynaptic8* was first described by Hernandez-Soriano (1973) in spontaneous semi-sterile mutants found in the cultivar Betzes (Hockett and Eslick, 1969). Two alleles *des8.k*, *des8.l* are known, both exhibiting similarly perturbed meiosis with univalents present at Metaphase I although the associated semi-sterility is reported to be stronger in *des8.l* (Figure 1a,b) (Hernandez-Soriano, 1973, Lundqvist et al., 1997). Here using a positional candidate-gene approach we show that *desynaptic8* mutants (Druka et al., 2010; Hernandes-Soriano, 1973) are associated with deletions in or of the barley homolog of *XRCC2* enabling us to compare the meiotic phenotype and effect on recombination of the mutants with that in the model plant Arabidopsis.

**Figure 1:**
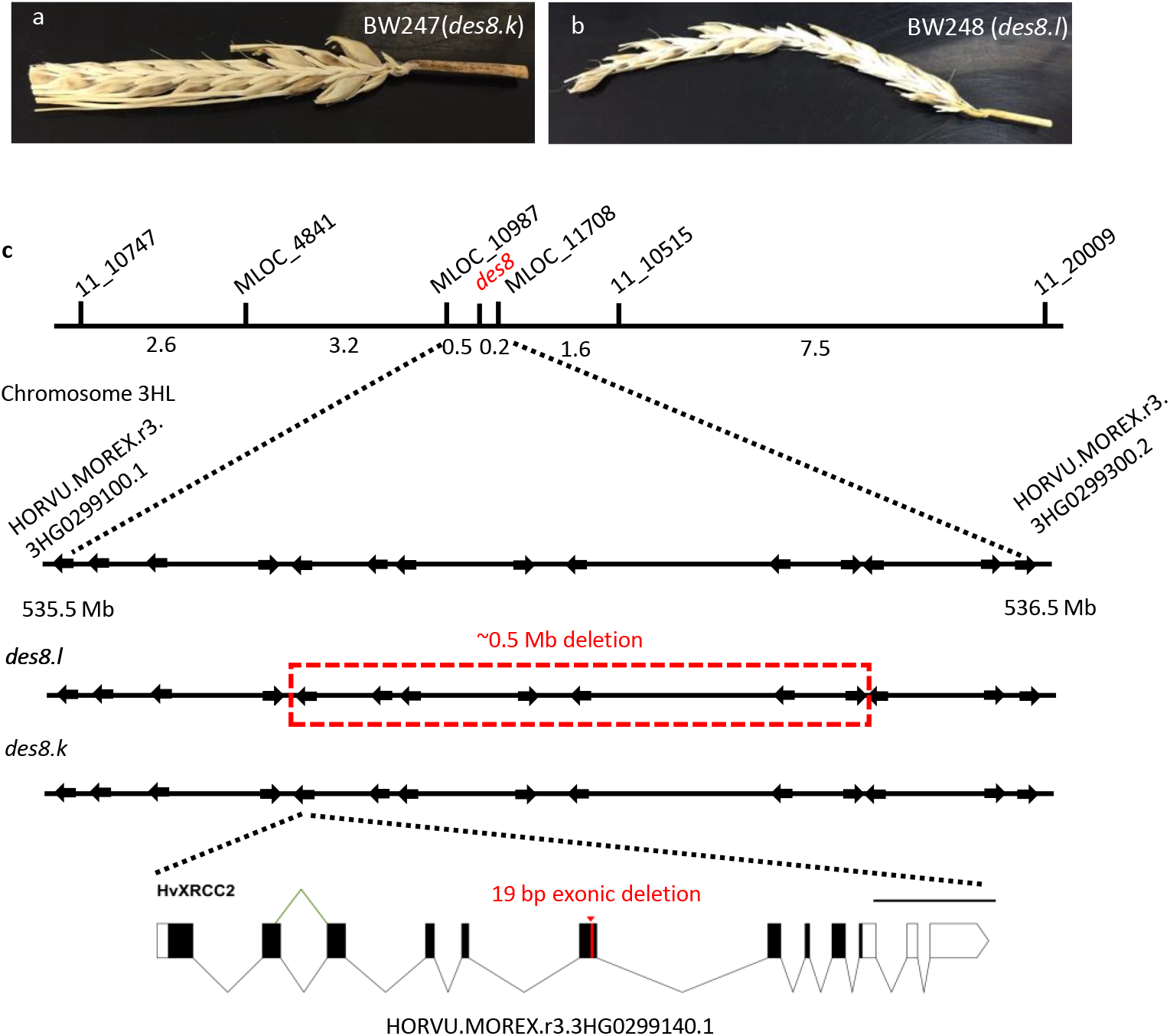
*des8* mapping. *desynaptic 8* mutants exhibit a semi-sterile phenotype (a,b). The *des8* region (c) was initially delineated between two SNP markers (11_20659 and 11_10515) on chromosome 3H and then fine mapped on an extended F_2_ population to a less than 0.5 cM region between two markers (MLOC_10907 and MLOC_11708) corresponding to a region between HORVU.MOREX.r3.3HG0299100.1 and HORVU.MOREX.r3.3HG0299300.2. *des8.l* in BW248 is a large deletion of 0.5Mb while *des8.k* in BW247 has a small 19bp exonic deletion in *HvXRCC2* (HORVU.MOREX.r3.3HG0299140)

## Materials and methods

### Plant and material preparation

The cultivar Bowman (wild-type; WT) and its near-isogenic lines BW247 (BC_2_F_3_; *des8.k*) and BW248 (BC_4_F_3_; *des8.l*) were grown under 16h of light at 18-20°C and 8h of dark at 16°C in general-purpose mix compost containing peat, sand, limestone, perlite, Celcote, Osmocote®, and Exemptor® was used in all the experiments. After 7 weeks from sowing, anthers were checked for meiotic stage using aceto-carmine spreads and fixed for cytology. For mapping and recombination studies, F_2_ and derived F_3_ populations resulting from BW247 (*des8.k*) x cv. Morex, BW248 (*des8.l*) x cv. Barke or BW248 (*des8.l*) x cv. Morex were grown in a glasshouse and young leaf tissue from (approx. 2 week-old seedlings) were collected into 96 well plates for DNA extraction as described previously (Arrieta et al., 2021). Plants were grown to maturity to allow for the assessment of fertility and harvest.

### Mapping and sequencing

Initial genetic mapping was based on 94 F_2_ individuals derived from a cross between BW248 (*des8.l*) and the cv Barke (WT) using segregation of the semi-sterile phenotype of *des8.l* as a Mendelian trait. Genotyping was carried out using an Illumina 384 SNP BeadXpress platform as previously (Colas et al., 2016). Subsequent mapping was carried out in 920 F_2_ individuals derived from a BW428 (*des8.l*) x Morex cross via the iterative development of custom KASPar© SNP assays (KBioscience). These Diagnostic KASP assays were designed as previously described (Arrieta et al., 2021) from alignments of genic sequences known to map in the genomic interval mined for polymorphism between BW248 and Morex. Initial genetic mapping of the BeadXpress data of the F_2_ segregating population, used segregation of the semi-sterile phenotype of *des8.l* as a Mendelian trait. JoinMap 4.0 (Kyazma) software was used to assign marker loci to linkage groups and two rounds of regression mapping used to order the loci within groups. For fine mapping, iterative rounds of SNP marker development were utilised on progressively smaller subsets of the F_2_ population shown to contain informative recombination events. Sequencing the coding domains of high confidence gene models was carried out by standard Sangar sequencing, as was confirmation of the BW247 (*des8.k*) cDNA sequence as previously described (Colas et al., 2016).

### Recombination study

The effect of *des8.l* on the recombination landscape was studied using the segregation of SNP loci in F_3_ families derived from specific F_2_ individuals from a BW248 (*des8.l*) x Barke cross that were homozygous for either the WT or mutant allele at *HvXRCC2*. Genotyping was carried out using the Barley 50K iSelect SNP array (Bayer et al., 2017) as previously described (Arrieta et al., 2021). Subsequent 50K data handling and recombination analysis were carried out using custom R scripts (see Data Availability). Briefly, monomorphic markers between BW248 and Barke genotypes were removed. F_3_ marker calls were converted to a numeric matrix representing a match to either parental genotype (0 and 2) or a heterozygous call (1). A sliding window was used to adjust genotyping calls to the median within a 20-marker window to account for poorly mapped markers. Outlier markers and F_3_ individuals containing a large number of expected transitions (>95% quantile and >98.5% quantile respectively) along the chromosome were removed. Informative (F_2_ heterozygous) and uninformative (F_2_ homozygous) regions were determined by the presence or absence of large monomorphic blocks matching either genotype in the original cross in F_3_ individuals grouped by F_2_ parent. Crossovers were counted at the junction between three continuous matches/mismatches to the F_2_ parental genotype in each direction. Recombination frequency for each marker for each segregating phenotype was calculated as the total number of crossovers observed divided by the number of observations for this marker within an F_2_ heterozygous region.

### Bleomycin/mitomycin-c DNA damage repair assay

Seeds of Bowman, BW247, and BW248 were sterilized with 70% EtOH for 30 seconds and rinsed 3 times for 5 minutes in with Sterile water. Seeds were subsequently surface sterilized with 7% Sodium Hypochlorite for 4 minutes and rinsed 3 times for 5 minutes, followed by 1 wash for 1 hour in sterile water. Seed were grown on 0.5% DUCHEFA Phyto agar (Melford Laboratories Ltd, UK) for the control containing Mitomycin C (Abcam, UK), 10mg/l and Bleomycin (Cambridge Bioscience, UK) 50mg/l respectively for 7 days at 21 degrees and the root length measured.

### DNA in situ hybridization

Anthers were fixed in Ethanol/Acetic acid (3:1) for 24 hours and stored in 70% Ethanol at 4°C until use. Slide preparation and DNA *in situ* hybridizations were performed as previously described (Colas et al., 2016; Higgins et al., 2012) using rDNA 5s-digoxigenin and rDNA 45s-biotin, barley sub-telomeric HvT01 FITC-dUTP and centromeric BAC7 Biotin-dUTP (Jasencakova et al., 2001).

### Immunocytology

Anthers were collected in working buffer (1X PBS / 0.5% Triton^TM^ X-100 / 0.5% Tween20) until fixation in 4% formaldehyde (in working buffer) for 20 minutes. Following 2 washes of 10 minutes in 1XPBS, anthers were tapped to release the meiocytes. The meiocyte suspensions (30µl) were transferred onto a *Polysine® slide (Poly-L-Lysine coated slides)* and left to air dry at room temperature. Slides were first blocked for 20 minutes in 5% Goat/Donkey serum prepared in working buffer, followed by an incubation in the primary antibody solution which consisted of one or multiple antibodies diluted in blocking solution in a wet chamber for 1 hour at room temperature. This was followed by 24-36h incubation at 4°C. The following antibodies were used: Rabbit TaASY1 (1:2000); Rat HvZYP1 (1:500); Guinea Pig HvDMC1 (1:200); Rat HvRAD51 (1:200), Rabbit HvMLH3 (1:500) and Rabbit H3K27me3; (Baker et al., 2015; Colas et al., 2016; Colas et al., 2019). HvRAD51 polyclonal antibody was developed for this study by immunizing a rat with 2 custom designed peptides as per the supplier’s protocol (Dundee Cell Product, UK). Slides were warmed for 30 minutes to 1 hour at room temperature before washing for 15 minutes in 1XPBS and incubating for up to 2 hours at room temperature in a secondary antibody solution. Secondary antibodies consisted of a mixture of anti-rabbit Alexa Fluor® (488, 568, or 647), anti-rat Alexa Fluor® (568, 488, or 647) and/or anti-Guinea Pig Alexa Fluor® (568 or 488) (Invitrogen^TM^) diluted in blocking solution (1:300). Slides were washed in 1xPBS, counterstained for 10 minutes in a wet chamber with Hoechst 33342 (2µg/ml, Invitrogen^TM^) and mounted in ProLong™ Gold antifade mounting medium (Thermo Fisher Scientific).

### Time course

We used a Click-iT™ EdU Alexa Fluor™ 647HCS assay kit (Invitrogen^TM^) as previously described (Colas et al., 2016). Fixed anthers were prepared for immuno-detection as described above with TaASY1 and HvDMC1 or HvRAD51, immediately followed by EdU detection as per the supplier’s protocol. EdU was detected after 45 minutes incubation instead of 30 minutes in the supplied protocol. Slides were counterstained with Hoechst 33342 (2µg/ml, Invitrogen^TM^) and mounted in ProLong™ Gold Antifade mounting medium (Thermo Fisher Scientific).

### Microscopy and Imaging

3D Confocal stack images (512 x 512, 12 bits) were acquired with LSM-Zeiss 710 using laser light 405, 488, 561, and 633nm sequentially. Projections of 3D pictures and light brightness/contrast adjustment were performed with Imaris 9.5.1 (Bitplane). 3D-SIM images were acquired on a DeltaVision OMX Blaze (GE Healthcare) for Laser light 405, 488, and 564nm as previously described (Colas et al., 2016). Image analyses were performed with imaris 9.5 as previously described (Colas et al., 2016) and Imaris image deconvolution clearview 9.5.

## Results

### Genetic mapping of <u>des8</u>

Both *des8.k* and *des8.l* have been used as donor parents in an extensive backcrossing programme using the cultivar Bowman as the recurrent parent generating the near isogenic lines BW247 (Bc_2_F_3_) and BW248 (Bc_4_F_3_) respectively (Druka et al., 2011). Initial genotyping of BW247 and BW248 with 3,072 SNPs (BOPA1 and BOPA2, Close et al., 2009) allowed comparisons of the NILs with the donor and recurrent parent cultivars and indicated the presence of introgressions on 3HL and 5HL for BW247 and 3H and 6HL in BW248 (Druka et al., 2011).

We initially mapped *des8.l* genetically as a Mendelian trait based on segregation of its semi-sterile phenotype in 94 F_2_ individuals derived from a cross between BW248 (*des8.l*) and the cv. Barke (WT) (Figure 1b). The use of an Illumina 384 SNP BeadXpress platform enabled *des8* to be mapped to the long arm of chromosome 3H between SNPs 11_10747 and 11_20009 which map within the genes HORVU.MOREX.r3.3HG0297410.1 and HORVU.MOREX.r3.3HG0302040.1 respectively. This position overlaps with the introgressions observed on chromosome 3H in BW247 and BW248 (Druka et al., 2011). Given a lack of polymorphism in the cross with Barke, subsequent mapping was carried out using 920 F_2_ individuals derived from a BW248 (*des8.l*) x cv. Morex cross. The iterative development of custom KASPar© SNP assays (KBioscience) derived from alignments of genic sequences known to map in this interval, were mined for polymorphism between BW248 and Morex, and these were used to delineate the interval containing *des8.l* to that bounded by MLOC_10987 (HORVU.MOREX.r3.3HG0299100.1) and MLOC_11708 (HORVU.MOREX.r3.3HG0299300.2) (Figure 1c).

The interval containing *des8.l* was thus mapped to within a single 1.37Mb BAC contig (contig_1227) (http://mips.helmholtz-muenchen.de/plant/barley/fpc/index.jsp) to a region containing 14 high confidence annotated genes (Figure 1c). Using the sequence information within BAC contig_1227, primers were designed across the coding domains of the genes bounded by flanking SNPs. This indicated that seven high confidence genes were missing in BW248 (*des8.l*) relative to Morex (HORVU.MOREX.r3.3HG0299140.1 to HORVU.MOREX.r3.3HG0299250.1), a deletion of over 474 kb (Figure 1c). Resequencing these seven genes in BW247 (*des8.k*) found that all were identical to the donor Betzes except for HORVU.MOREX.r3.3HG0299140.1 which carried an 19bp exonic deletion (Figure 1c). These results indicate that *desynaptic-8* mutants are caused by mutations involving HORVU.MOREX.r3.3HG0299140.1, the barley homolog of X-Ray Repair Cross Complementing 2, XRCC2.

*HvXRRC2* is comprised of 12 exons, contains two 3’UTR introns (Figure 1c) and encodes a protein of 347 amino acids that shows a high level of conservation to other known XRRC2 proteins, in particular over the P-Loop_NTPase superfamily domain that contains the walker A and walker B motifs (Figure S1). The 19 bp deletion in the *des8.k* allele (BW247) (Figure S2) was confirmed by cDNA sequencing and is in the sixth exon of *HvXRCC2*. It is predicted to produce a truncated protein of 262 amino acids (Figure 1c and Figure S2). An expression atlas across 16 tissues of cv. Morex (Rapazote-Flores et al., 2019) indicated that *HvXRCC2* is expressed in all tissues but highly expressed in developing inflorescence (Figure S3a). In addition, a meiotic specific expression dataset from the cultivar Golden Promise indicated that *HvXRCC2* was highly expressed in meiocytes during G2 and leptotene/zygotene stages but less expressed from pachytene onward (Barakate et al., 2021) (Figure S3b). *HvXRCC2* exists in two main isoforms, BAnTr.GP.3HG012678.1, expression of which appears to be meiosis specific, and BAnTr.GP.3HG012678.2 which has an alternative 5’ splice site of 3 nucleotides at the end of exon 2 but is expressed at a lower level than isoform 1 (Figure S3b). The presence of these two isoforms is supported in the BaRTv1.0 barley transcript dataset visualised via the EoRNA database (Milne et al., 2021). The latter transcript (BaRT1_0u23042.002) potentially has a more mitotic role with stronger expression in apical meristem libraries, although interpretation is complicated as all *HvXRCC2* transcripts in EoRNA are 5’ truncated possibly due to issues in earlier genome coverage (Mascher et al., 2017) and resultant gene models (e.g. HORVU3Hr1G082620.28).

### HvXRCC2 has a somatic role in barley

Given the apparent ubiquitous expression of XRCC2 in the EoRNA barley expression database (https://ics.hutton.ac.uk/eorna/index.html) and the report of an effect on both mitosis and meiosis in Arabidopsis (Da Ines et al., 2013a; Wang et al., 2014) the *desynaptic8* mutants were investigated for a possible somatic role. We tested the response of the *des8* lines to the application of genotoxic agents using the DNA cross-linking agent Mitomycin C and radiomimetic agent Bleomycin (Da Ines et al., 2013a; Liu and Lim 2005; Wang et al., 2014). The root lengths of the NILs containing *des8* alleles were not significantly different to those of the Bowman WT in the absence of genotoxic agents and the presence of genotoxic agents had a significant effect on root length (p<0.005, Welch’s t-test) in all lines. No significant difference was found between lines when grown in the presence of Bleomycin however in the presence of Mitomycin C, BW248 (*des8.l*) did have significantly shorter roots than Bowman WT (p<0.05, Welch’s t-test) although BW427 (*des8.k*) did not (Figure S4).

### des8.k and des8.l undergo complete synapsis despite a short delay at the beginning of the synaptonemal complex formation

Using TaASY1 and HvZYP1 antibodies, the progression of synapsis was followed in Bowman (WT), BW247 (*des8.k*), and BW248 (*des8.l*) meiotic cells (Figure 2). In all three lines, ZYP1 started to polymerize at one side of the nucleus during leptotene/zygotene (Figure 2a-f). ZYP1 polymerized in between the homologous chromosomes to form continuous ZYP1 filaments in the WT (Figure 2d), but not in BW247 and BW248 where ZYP1 formed short discontinuous stretches (Figure 2e-f) at the same stage. Later, during zygotene, ZYP1 polymerized along the chromosomes in all genotypes but the appearance of un-synapsed ASY1 loops was frequent in BW247 and BW248 (Figure 2h,i, arrows), suggesting that synapsis progression suffered some delays in the mutants (Colas et al., 2016). By pachytene, synapsis was completed as shown by the full ZYP1 polymerization and thick chromosome axes in all genotypes (Figure 2j-l). All three genotypes reached diplotene, exhibiting tinsel chromosome structures (Colas et al., 2017) (Figure 2m-o), though ZYP1 aggregates were visible on both mutants but not in the WT (Figure 2n,o, arrows).

**Figure 2:**
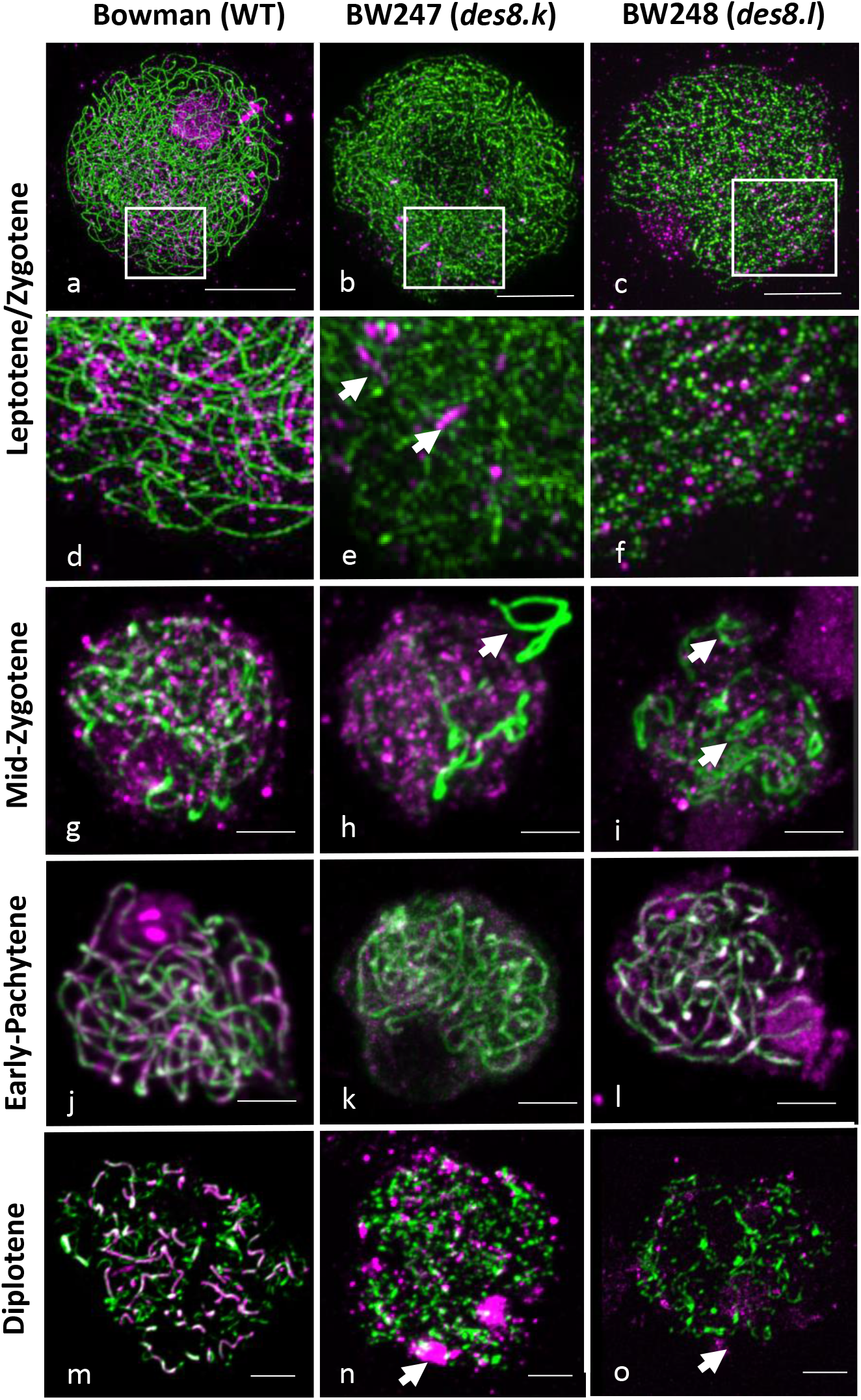
Comparison of synapsis in Bowman, BW247 and BW248. Synapsis of all genotypes tracked by ASY1 (green) and ZYP1 (magenta) at leptotene/zygotene (a-f), mid-zygotene (g-i), pachytene (j-l) and diplotene stages (m-o). Arrows in (e) indicate ZYP1 stretches and arrows in (h) and (i), indicate un-synapsed ASY1 labelled chromosomes. Scale bar 5 µm.

### Synapsis delay affects RAD51 and DMC1’s behaviour in both des8.k *and* des8.l

Due to the presence of the atypical short ZYP1 stretches in BW247 and BW248 during leptotene/early zygotene, and the known involvement of XRCC2 in replication fork progression (Liu and Lim, 2005; Saxena et al., 2018) we hypothesized that entry into meiosis could be compromised in the *des8* mutants. We conducted a time course experiment to monitor the start of synapsis by injecting 1-1.5 cm long spikes with an EDU solution for 2 hours and collecting the injected spikes at 24 and 40 hours as previously described (Colas et al., 2016). Cells at leptotene/zygotene were checked for EDU, ASY1 axis and the early recombination proteins HvDMC1 and HvRAD51 (Figure 3). As cells only start meiosis when replication is completed any EDU labelling indicated that the cells had been replicating in the presence of EDU within the time of incubation.

**Figure 3:**
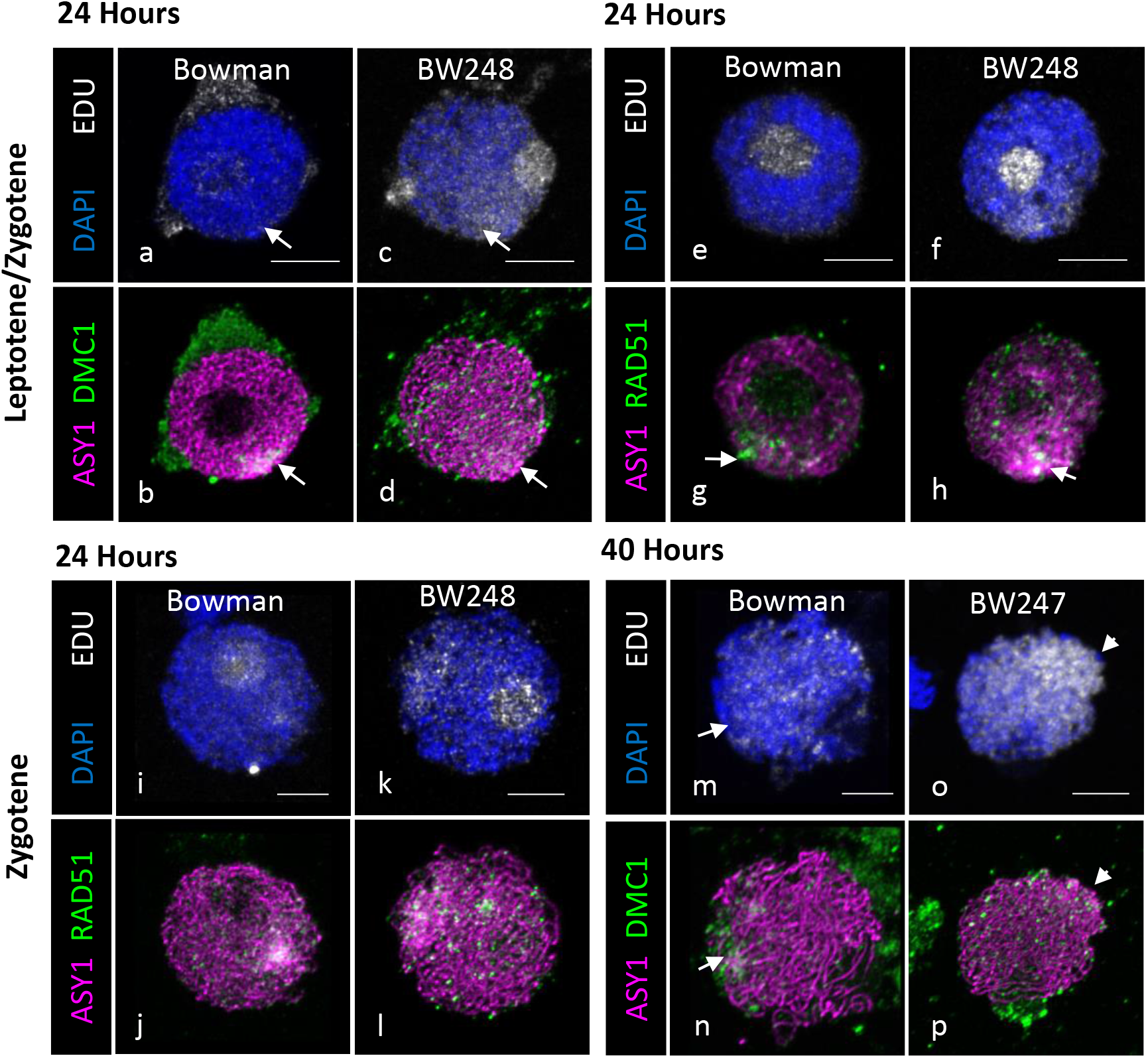
Time course analysis of early recombination events. Time course of Bowman and *des8* mutants using EDU (in white), ASY1 (in magenta) and HvDMC1 (in green) or HvRAD51 (in green). Labelling at 24 hours with HvDMC1 (b,d) and HvRAD51 (g,h) in cells at leptotene/zygotene in Bowman (a,b,e,f) and BW248 (c,d,f,h). Labelling at 24 hours with HvRAD51 in cells at zygotene in Bowman (i,j) and BW248 (k,l). Labelling at 40 hours with HvDMC1 in cells at zygotene in Bowman (m,n) and BW248 (o,p). Arrows indicates the telomere clustering position. Scale bar 5 µm.

After 24 hours, leptotene/zygotene cells in Bowman were not EDU labelled, indicating replication was completed in these cells prior to EDU injection (Figure 3a and 3e). At this stage ASY1 axes are fully formed and both HvDMC1 and HvRAD51 foci congregate towards a brighter ASY1 region that resembles the telomere cluster (Figure 3b and 3g arrow). BW248 cells at the same stage were partially labelled with EDU, indicating that some replication had occurred within the last 24 hours (Figure 3c and 3f). As in the WT, ASY1 axes appeared fully formed and both HvDMC1 and HvRAD51 foci congregated towards one side of the nucleus (Figure 3d and 3h arrow). HvDMC1 foci were however more dispersed within BW248 nuclei compared to the WT (Figure 3d).

In WT cells that were at zygotene at 24 hours, faint EDU labelling was evident with a brighter spot at one side of the nucleus (Figure 3i) indicating that these cells had recently finished late replication. ASY1 axes were linear and RAD51 foci clustered to the EDU labelled side of the nuclei (Figure 3j). We observed a similar result in BW248, but with stronger EDU labelling than in WT (Figure 3k) indicating that replication was slightly longer in the mutant. RAD51 foci were also more dispersed within the nuclei compared to WT (Figure 3l).

At 40 hours, most labelled WT nuclei were at zygotene and were homogenously labelled with EDU (Figure 3m). The WT ASY1 axes were thick but were beginning to be less evident in stretches where synapsis had been initiated. Similarly, HvDMC1 foci were still concentrated to one side of the nucleus but foci were showing signs of starting to diffuse (Figure 3). BW247 nuclei were also labelled with EDU but the signal intensity was much brighter than in the WT with an even stronger signal at one side of the nucleus (Figure 3o). Similar to the WT, HvDMC1 foci were still concentrated at one side of the nucleus in BW247 though starting to diffuse (Figure 3p). However, the clusters of HvDMC1 foci in BW247 and BW248 were not as tight as in WT at any time point (Figure 3d, 3p).

The time course experiment suggested that replication was affected, taking longer in both mutants, with more EDU signal in the mutants than WT at comparable stages. In addition to the delay in replication the mutant lines exhibited an altered behaviour of DMC1 and RAD51 foci compared to Bowman. To confirm this, we followed the behaviour of HvRAD51 and HvDMC1 during synapsis using both TaASY1 and HvZYP1 antibodies (Figure 4 and Figure 5). We confirmed that in WT during leptotene HvDMC1 foci congregated towards one side of the nucleus, where the bouquet normally forms (Figure 4a_1,2_). This HvDMC1 localization is maintained at early zygotene (Figure 4b_1,2_) and by mid-zygotene, HvDMC1 foci are dispersed within the nucleus although some polarity is still retained around the ZYP1 axes (Figure 4c_1,2_). In the *des8* mutants, HvDMC1 foci were present but more diffuse in the nucleus and appeared to have lost their polarity at leptotene (Figure 4d_1,2_ and Figure 4g_1,2_) and zygotene stages (Figure 4e_1,2_ and Figure 4h_1,2_). By mid-zygotene, HvDMC1 still appeared more diffuse in the mutants compared to the WT (Figure 4f_1,2_ and Figure 4i_1,2_).

**Figure 4:**
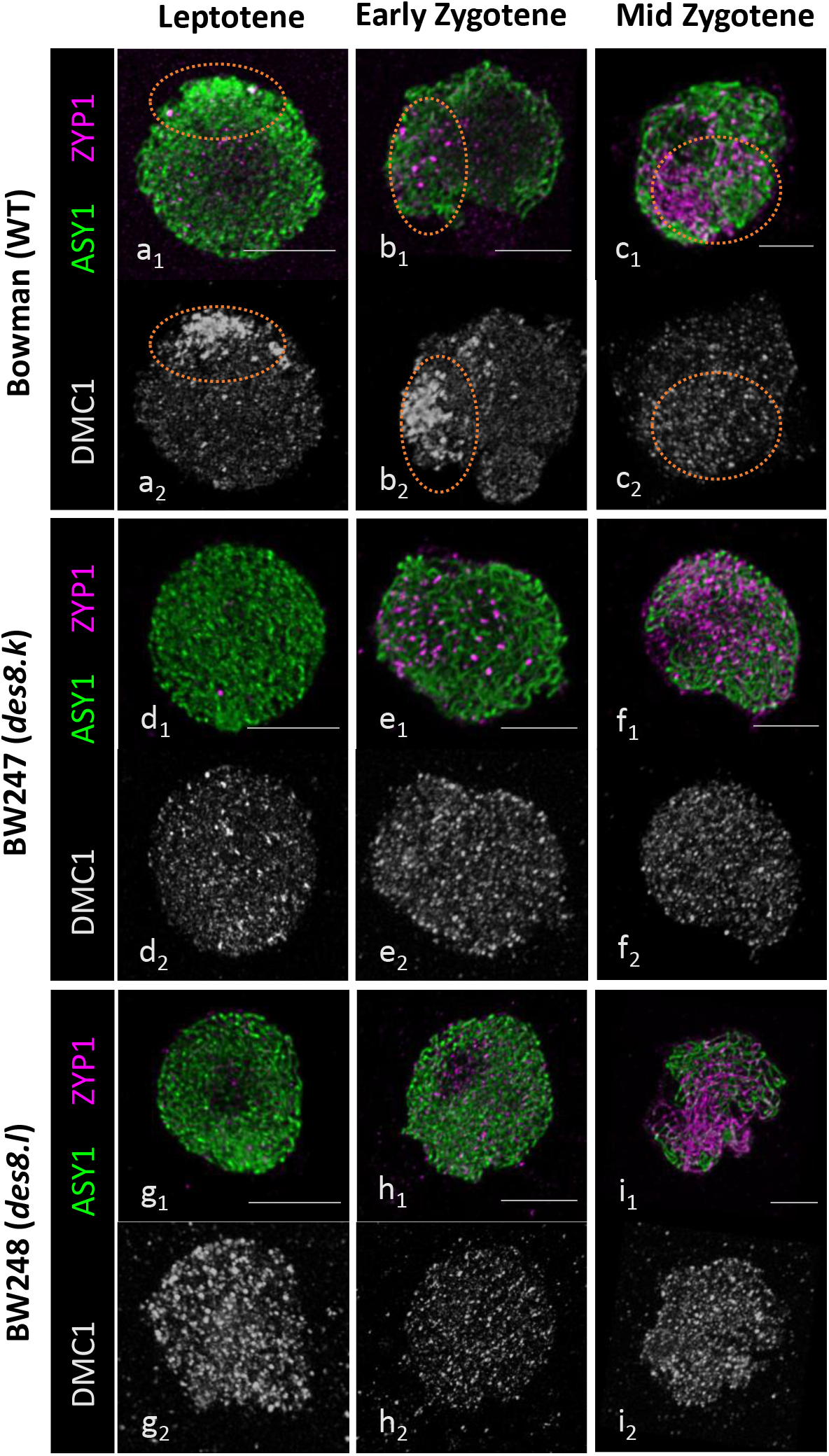
DMC1 behaviour during synapsis. Immunolabelling of Bowman (a-c), BW247 (d-f) and BW248 (g-i) with ASY1 (green), ZYP1 (magenta) and DMC1 (white). Orange ellipses indicate the telomere area where DMC1 foci normally start loading.

**Figure 5:**
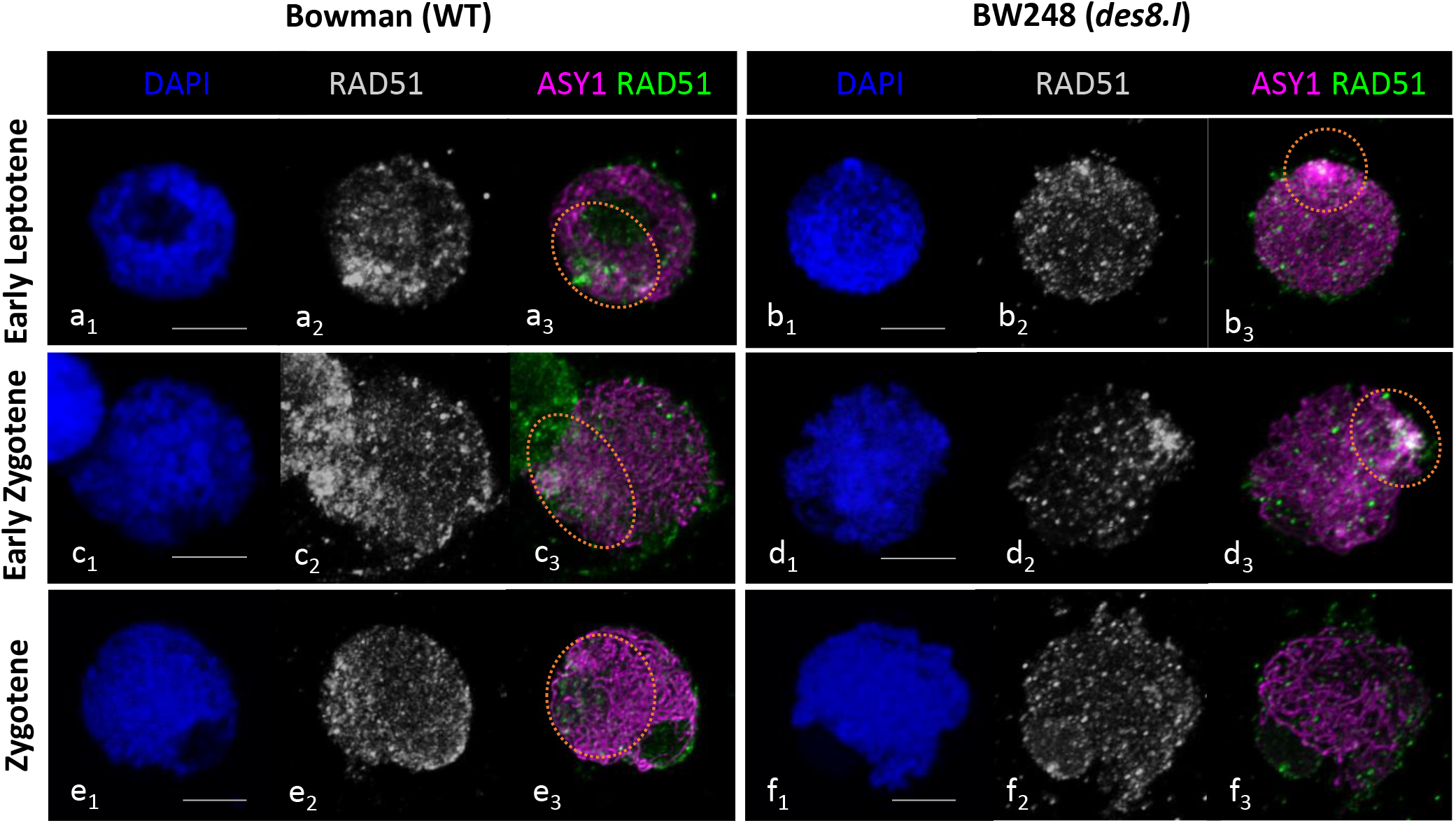
RAD51 behaviour during synapsis. Immunolabelling of Bowman (a-c), and BW248 (d-f) with ASY1 (magenta) and RAD51 (green/white). Orange ellipses indicate the telomere area where RAD51 foci normally start loading. Scale bar 5 µm

Similarly, HvRAD51 foci were also present in BW248 (Figure 5), but unlike HvDMC1, they retained some polarity during early leptotene in both WT (Figure 5a_1-3_) and the mutant (Figure 5b_1-3_). This HvRAD51 polarity was maintained at early zygotene in both WT (Figure 5c_1-3_) and the mutant (Figure 5d_1-3_). At later zygotene, HvRAD51 retained some polarity in the WT (Figure 5e_1-3_), but foci were more diffuse in the mutant (Figure 5f_1-3_). Attempts were made to determine if the number of HvDMC1 and HvRAD51 foci was also altered in the mutants, but high-count standard deviations meant that no significant differences were found (Figure S5).

### des8.k and des8.l both exhibit reduced cross-overs

At Metaphase I, Bowman showed seven ring bivalents with an average of 14.9 ±3 (n=50) chiasmata / cell (Figure 6a and Figure S6A). BW247 and BW248 displayed abnormal metaphase with an average of 8.9 (±1.5 n=20) and 8.8 (±2.4 n=36) chiasmata respectively (Figure 6b,c and Figure S6A) representing a 40% reduction compared to WT. The metaphase spreads also showed that, as previously reported in the original mutants, both BW247 and BW248 exhibit a range of ring-and rod bivalents and univalents, suggesting that some of the obligate crossovers had been lost (Figure S7). We used fluorescence *in situ* hybridization (FISH) with probes against 45S and 5S rDNA to determine chiasma frequencies of individual barley chromosomes of Bowman and BW248 and found that chiasma number was reduced in all seven barley chromosomes in the mutant, particularly on chromosomes 4H and 6H (Figure S8). During anaphase I, the centromeres pulled the homologous chromosomes cleanly to each side of the nucleus in Bowman (Figure 6d) but both BW247 and BW248 showed lagging chromosomes, chromosome bridges (Figure 6e,f, white arrows) and abnormal chromosome orientation (Figure 6f, white star). Although finding pachytene cells was challenging, a comparison of MLH3 labelled foci marking class I CO was carried out in all genotypes. We found 7.8 (±1.9, n=5), 7.1 (±1.4, n=20), and 17.2 (± 2.3, n=14) MLH3 foci in BW247, BW248, and Bowman respectively again indicating a significant difference between the WT and two mutant *des8* lines (Figure 6g,h and Figure S6B).

**Figure 6:**
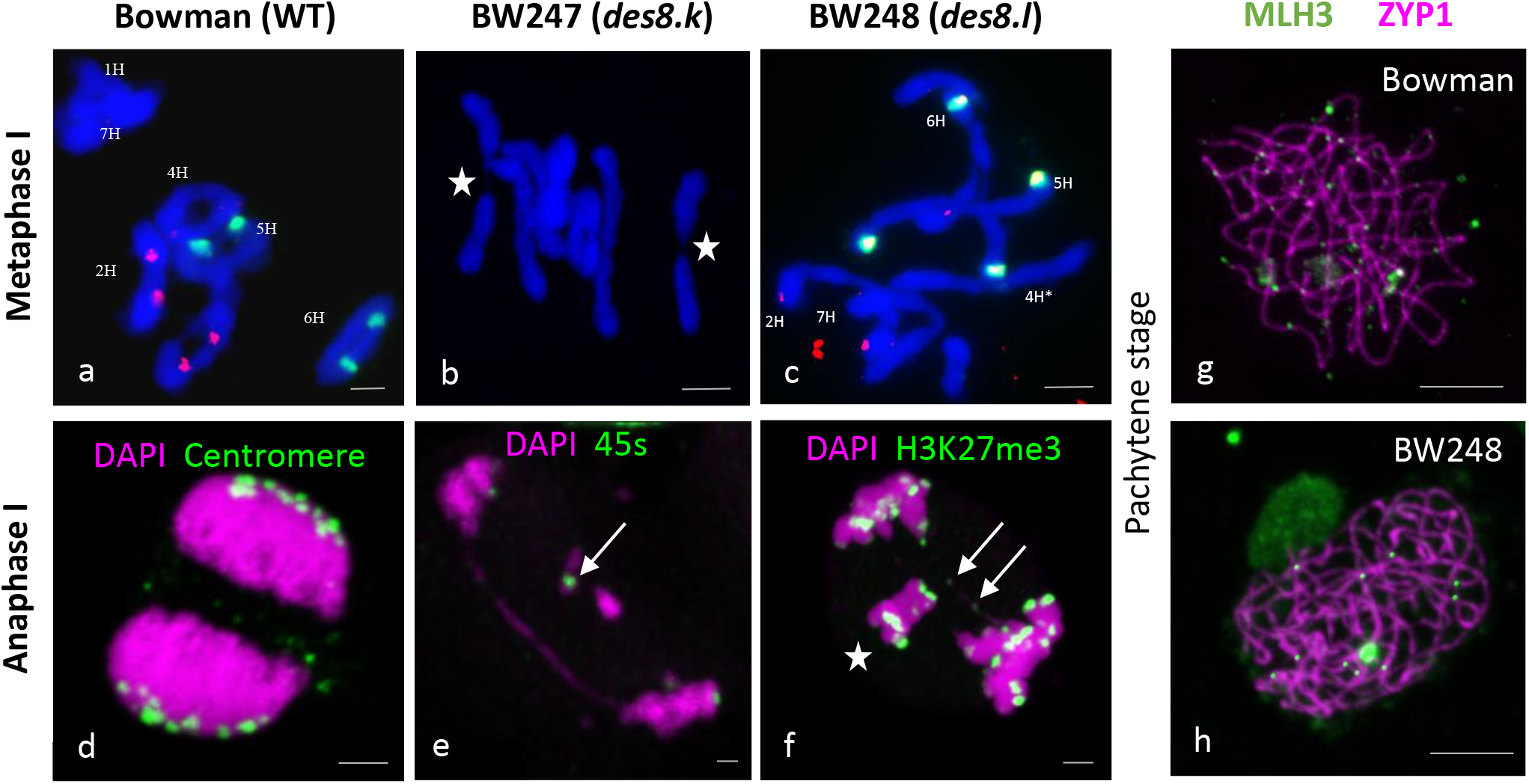
Metaphase and chromosome segregation. Metaphase of Bowman (a), BW247 (b) and BW248 (c). Bowman and BW248 metaphase spreads were labelled with 45s (red) and 5s (green) probes. Chromosome segregation at Anaphase I in Bowman (d), BW247 (e) and BW248 (f) with chromatin labelled in magenta and centromeres, 45s or H3K27me3 labelled in green. Pachytene cells of Bowman (g) and BW248 (h) were labelled with HvZYP1 (magenta) and HvMLH3 (green). Scale bar 5µm

### des8.l exhibits reduced recombination in genetic mapping

To study the change in recombination, F_3_ individuals from the BW248 (*des8.l*) x Morex cross were genotyped with the Barley 50K iSelect SNP array (Bayer et al., 2017). Our analysis utilized data from 11916 polymorphic SNP loci on 82 F_3_ individuals from 10 homozygous *des8* WT F_2_ families and 78 F_3_ individuals from 6 homozygous *des8.l* F_2_ families. Across all chromosomes, there was a reduction in interstitial recombination frequency in *des8.l* compared to WT (Figure 7). The average number of crossovers observed was generally reduced across all chromosomes (16% to 66% reduction; Table S1) with the exception of chromosome 3H, which showed almost no difference. This reduction in recombination reached statistical significance (T-test with Benjamini Hochberg correction; Table S1) in chromosomes 2H (p<0.0001), 4H (p<0.01), 5H (p<0.0001), and 6H (p=0.02) but not in chromosomes 1H (p=0.066) and 7H (p=0.31) (Figure S9).

**Figure 7:**
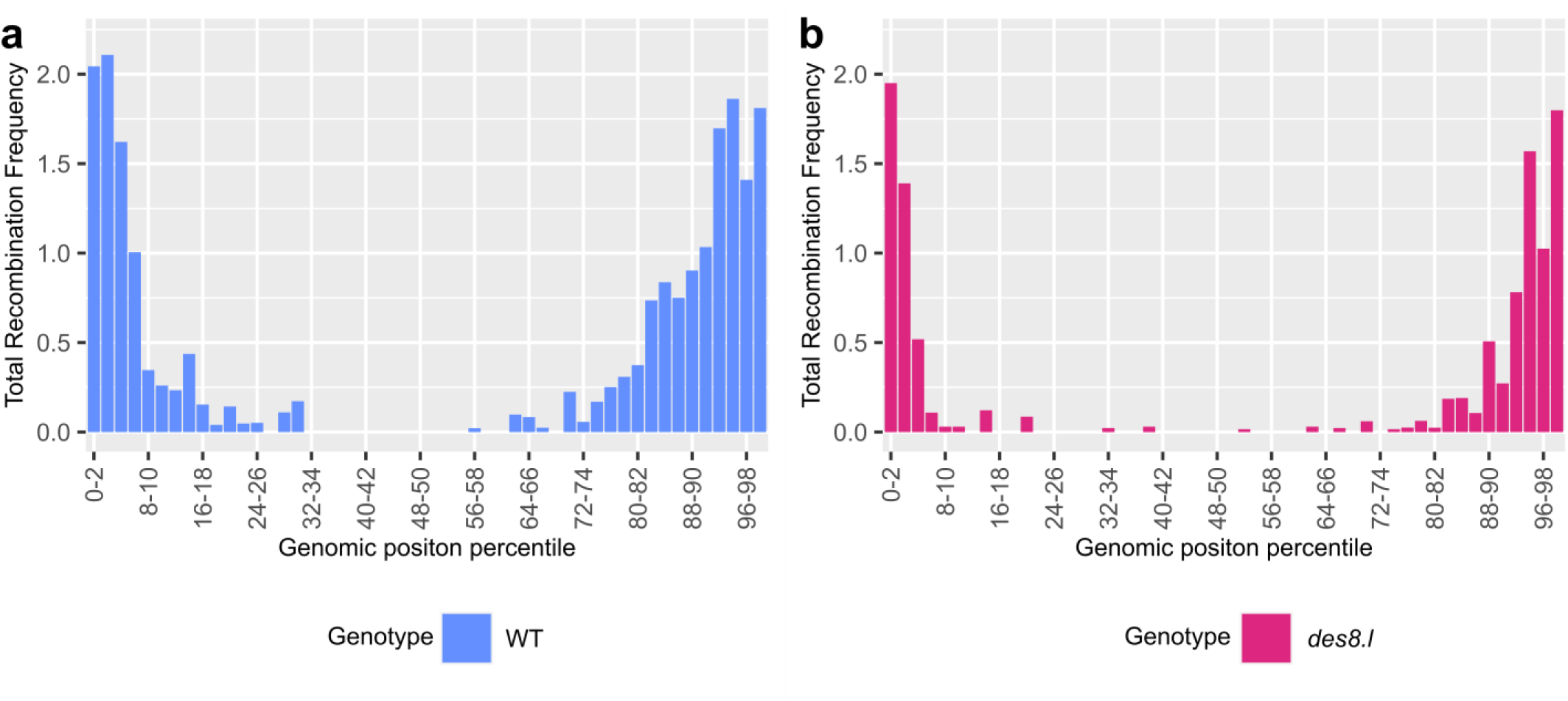
Genetic map length comparisons of F_3_ families with WT and *des8.l*. Recombination found in F_3_ families derived from F_2_ individuals homozygous for either WT (a, Bowman) (in blue) or *des8.l* (b, BW248) (in pink) at *HvXRCC2*.

## Discussion

This study has shown that the semi-sterile *desynaptic8* mutants in barley are associated with deletions in or of the barley homolog of *XRCC2* (*HvXRCC2*). Effective null *xrcc2* mutants in the Chinese hamster cell line irs1 are sensitive to genotoxic agents and display severe meiotic phenotypes with chromosome mis-segregation (Mozdarani et al., 2001). An *XRCC2* point mutation in humans and mice maintains high expression in testes but results in meiotic arrest and infertility (Yang et al., 2018). The potential sensitivity of *des8.l* to Mitomycin C shown here indicates that *XRCC2* is likely to have a conserved role in DNA repair in barley, consistent with observations in both mammals and plants (Liu and Lim, 2005; Wang et al., 2014).

The two described *desynaptic8* alleles identified as spontaneous semi-sterile mutants were originally found in the cultivar Betzes (Hockett and Eslick, 1969) with both alleles (*des8.k*, *des8.l)* exhibiting similarly perturbed meiosis with univalents present at Metaphase I (Hernandez-Soriano, 1973). We found that these *Hvxrcc2* mutants initially exhibited normal meiotic progression, albeit with a small delay in initiation, with completion of synapsis. However, the absence of *HvXRCC2* subsequently led to a dramatic reduction of crossovers, chromosome mis-segregation and infertility, suggesting that *HvXRCC2* has a major role in recombination in barley. The mutant phenotype of both alleles was very similar, with the potentially truncated allele *des8.k* showing similar disruption to meiotic progression (indeed potentially displaying a greater delay) and ultimately the same level of reduction of CO as the knock out allele *des8.l*. The similarity of the phenotypes of the two *des8* mutants (truncated and KO) highlights the potential importance of the C-terminal domain in the protein structure and in the interaction of Rad51 paralogs within the BCDX2 complex (Miller et al., 2004).

Both *desynaptic8* mutants exhibited a delay at the entry of meiosis, potentially related to XRCC2’s involvement in a genome ‘caretaker’ role and the resolution of DNA replication fork dynamics (Liu and Lim, 2005). Both *desynaptic8* mutants exhibited a delay in the initiation of synapsis and a noticeably more diffuse clustering of DMC1 foci compared to WT. This meiotic phenotype may relate to the known delay in (but not an absence of) RAD51 foci formation in *XRCC2* mutant hamster cells (Liu, 2002). The more diffuse DMC1 clustering may possibly reflect an issue in the recruitment of RAD51 and DMC1 in a timely manner (Da Ines et al., 2013b; Pradillo et al., 2012). Although differences in timing and distribution were noticed, no significant differences in DMC1 foci number were found, though this may be due to the inherent difficulties in such estimates. It is therefore possible that final DMC1 foci number is maintained in the *desynaptic8* mutants but that the dynamics of recruitment and retention may be compromised.

The observed reduction in crossover counts at metaphase I and in recombination frequency shown in the genetic mapping of the *desynaptic8* mutants relative to WT could relate directly to the known role of XRCC2 in homologous recombination (Liu et al., 1998, Liu, 2002; Masson et al., 2001). However, the distal distribution of chiasmata observed cytologically in the *desynaptic8* mutants and the reduction in proximal recombination found through genetic mapping may also correspond to the observed delay at the beginning of meiosis and synapsis initiation. Such a change in distribution would align with the observed delay in synapsis and a subsequent disconnect with the spatiotemporal control of meiotic progression displayed by large genome cereals such as barley (Higgins et al., 2012). It is likely that the skewed distribution of recombination found with the genetic mapping is potentially an underestimate of the effect of the *desynaptic8* mutant alleles at *HvXRCC2* given the mapping inherently selects against gametes resulting from chromosome mis-segregation that was evident cytologically. In addition, the chiasmata counts from Metaphase I spreads may underestimate CO numbers in WT as discussed previously (Colas et al., 2016), but are congruent with the MLH3 foci comparison between the mutants and the WT. Both experiments found an overall reduction in CO/recombination frequency with inter-chromosomal variation in the strength of this effect, though the reason for this variation in the effect of mutations in *HvXRCC2* is unclear.

The mutant phenotype displayed by *desynaptic8* barley plants is perhaps unexpected given the reported phenotypes shown by *XRCC2* mutants in other plant species. A recent study using VIGS induced knockdown of *XRCC2* in tetraploid wheat reported no associated change in the number of COs per chromosome but potentially some significant local effects in both pericentric and subtelomeric regions (Raz et al., 2021). A more striking difference is to the phenotype reported in Arabidopsis where the knock-out *xrcc2* mutant is fully fertile and exhibits normal chromosome pairing, synapsis, and correct chromosome segregation (Bleuyard et al., 2005; Bleuyard and White, 2004; Da Ines et al., 2013a). Moreover, it shows an increased CO rate and recombination (Da Ines et al., 2013a). Comparison between the two species is complicated by the different mutations underpinning the studies with the Arabidopsis work based on a T-DNA insertion mutant (*atxrcc2-1*) with the T-DNA inserted in intron 5, 3 bp after the end of exon 5, that produces a truncated mRNA (Bleuyard et al., 2005). The Arabidopsis allele (*atxrcc2-1*) is thus potentially analogous to the barley *des8.k* (BW247) mutant given the similar position of the truncation induced by the two mutation events (Supp Fig S1). However, the meiotic phenotypes of these two analogous mutants are very different, implying that the gene has different meiotic roles in the two species.

Despite the generally high conservation of the function of meiotic genes between organisms, significant differences in phenotype have previously been observed between plant species in meiotic mutant studies. For example, the effect of mutations in *MLH3* on synapsis in studies in barley compared to Arabidopsis (Colas et al., 2016; Jackson et al., 2006) and mutations in *ZYP1* homologs in rice compared to Arabidopsis or barley (Wang et al., 2010; Higgins et al., 2005; Barakate et al., 2014) are fundamentally different. Such comparsions are often complicated by the use of different mutagenesis strategies and the strength of the particular mutation events studied. However, the differences in mutant *XRCC2* phenotypes described here in barley compared to those previously described in Arabidopsis indicate a fundamental difference in the roles the protein plays in meiosis in the two species.

The mutant phenotype displayed by *desynaptic8* barley plants is similar to the phenotype exhibited in mammalian systems where *xrcc2* mutants are sterile, with clearly disturbed meiotic phenotypes alongside reduced recombination and aneuploidy (Cui et al., 1999; Griffin et al., 2000; Mozdarani et al., 2001). Again, the barley mutant phenotype for *xrcc2*, as is the case for *ZYP1* (Barakate et al., 2014) matches the expectation derived from non-plant models implying that, in these cases at least, model plant species with small genome sizes have the potential to display atypical mutant phenotypes reflecting non-canonical roles for some meiotic genes. Thus, barley displays mutant phenotypes that are largely aligned with expectations from other model eukaryotic systems and that any changes in patterns of recombination occur within the skewed distribution of recombination associated with the spatial and temporal control of meiotic progression displayed by large genome cereals (Higgins et al., 2012). The perturbed meiotic phenotype and reduction in recombination exhibited by the *desynaptic8* mutants in barley indicate that *XRCC2* is not a good gene candidate for practical exploitation to increase recombination frequency in crop plants. *XRCC2* is thus unlike *FANCM* or *RecQ4* which increase crossover counts in Arabidopsis and do show similar phenotypes in crop species including barley (Mieulet et al., 2018, Arrieta et al., 2021).

## Supplementary Data

**Figure S1:** Comparison of Arabidopsis and barley XRCC2 protein sequences and positions of studied mutant events.

**Figure S2:** Details of *des8.k* deletion in Exon 7 of *HvXRCC2*.

**Figure S3:** Expression levels of *HvXRCC2* shown in general and meiotic specific expression datasets.

**Figure S4**: Seedling root growth differences demonstrating Mitomycin and Bleomycin sensitivity

**Figure S5**: HvDMC1 and HvRAD51 foci counts at leptotene and zygotene for Bowman and *des8* mutants.

**Figure S6:** Chiasmata and MLH3 foci counts for Bowman and *des8* mutants demonstrating reduction in number of crossovers.

**Figure S7:** Range of Metaphase configurations shown by BW247 and BW248.

**Figure S8:** Comparison of mean chiasmata frequency for each chromosome in Bowman and BW248.

**Figure S9**: Comparison of recombination in F_3_ lines derived from F_2_ lines homozygous for either WT or *des8.I*.

**Table S1:** Comparison of crossover number for F_3_ lines derived from F_2_ lines homozygous for either WT *or des8.l*.

## Acknowledgements

We gratefully acknowledge the supervision of A.S. by Dr Eugenio Sanchez-Moran at the University of Birmingham and Professor Nils Stein and Dr. Sebastian Beier at the Leibniz Institute of Plant Genetics and Crop Plant Research (IPK) for the pre-publication access to Morex BAC contig sequence data.

## Authors contribution

IC, RW and LR: conceptualization; IC, LR, SJA and RW: methodology; IC, MA, MS and LR: formal analysis; IC, MM, AS, AA, MTK: investigation; IC, RW and LR: writing original draft; IC, MA, MS, JO, RW and LR: writing – review & editing; IC, MS and MA: visualisation; IC, SJA, LR and RW: funding acquisition.

## Data availability

The data are available upon request from the corresponding author. F_3_ recombination data are available on FigShare (DOI). Scripts used in 50K recombination analysis are available at github.com/BioJNO/des8.

